# Adaptation of the tetracycline-repressible system for modulating the expression of essential genes in *Cryptococcus neoformans*

**DOI:** 10.1101/2024.11.29.626100

**Authors:** Ci Fu, Nicole Robbins, Leah E. Cowen

## Abstract

The opportunistic human fungal pathogen *Cryptococcus neoformans* has an enormous impact on human health as the causative agent of cryptococcal meningitis, an AIDS-defining illness that results in significant human mortality. To combat *C. neoformans* infections, there is a dire need to expand our current antifungal arsenal. Essential gene products often serve as ideal targets for antimicrobials, and thus identifying and characterizing essential genes in a pathogen of interest is critical for drug development. Unfortunately, characterization of essential genes in *C. neoformans* is limited due to its haploid nature and lack of genetic tools for generating effective conditional-expression mutants. To date, the copper-repressible promoter *pCTR4* is the mostly widely used system to regulate essential gene expression, however, its expression is leaky and copper has pleiotropic effects. In diverse fungal species, including *Saccharomyces cerevisiae, Candida albicans*, and *Candida auris*, the tetracycline-repressible promoter system is a powerful tool to regulate gene expression, however, it has yet to be adapted for *C. neoformans*. In this study, we successfully implemented the tetracycline-repressible system in *C. neoformans* to regulate the expression of the essential gene *HSP90*. Supplementation of cultures with the tetracycline analog doxycycline efficiently depleted *HSP90* at both transcript and protein levels and inhibited *C. neoformans* growth and viability. Thus, this work unveils a novel approach to generate conditional-expression mutants in *C. neoformans*, providing unprecedented potential to systematically study essential gene function in this important human fungal pathogen.

## Importance

Invasive fungal infections cause millions of deaths annually, while the number of antifungals available to combat these pathogens is limited to only three classes: polyenes, azoles, and echinocandins. The largest source of novel antifungal drug targets are essential gene products, which are required for cellular viability. However, tools to identify and characterize essential genes in *C. neoformans* are extremely limited. Here, we adapted the tetracycline-repressible promoter system that has been widely used in other organisms to study essential gene function for use in *C. neoformans*. By placing this regulatable promoter upstream of the essential gene *HSP90*, we confirmed that growth of the strains in the presence of the tetracycline analog doxycycline results in depletion of essential gene expression. This approach provides a significant advance for the systematic study of essential genes in *C. neoformans*.

## Observation

Invasive fungal pathogens infect approximately 6.5 million people and directly contribute to 2.5 million deaths annually worldwide, making them a leading public health concern (1). Among the most problematic fungal pathogens, *Cryptococcus neoformans* affects 194 000 people and results in 147 000 deaths each year (1). Even with best available therapies, mortality rates remain high because the number of drug classes that have distinct targets in fungi is limited, and the utility of current antifungal drugs is compromised by either dose-limiting host toxicity or the frequent emergence of resistance (2). Thus, the discovery of new targets for therapeutic intervention is urgently needed.

Fungal-specific essential genes provide attractive drug targets as the majority of antimicrobials in clinical use target functions essential for pathogen viability, motivating systematic analysis of genes required for fungal survival. While genome-scale mutant collections that include coverage of essential genes are available in the model yeast *Saccharomyces cerevisiae* as well as the evolutionary divergent fungal pathogen *Candida albicans* (3-6), analogous tools in *C. neoformans* remain limited. *C. neoformans* deletion collections have been employed to further our understanding of its biology and vulnerabilities (7, 8), however, such resources only cover 62% of predicted protein-coding genes, and exclude essential genes (8). Further, tools available for characterizing essential genes in *C. neoformans* are limited. Galactose-inducible and copper-repressible promoters have been used to control *C. neoformans* gene expression (9, 10), however, leaky expression, interference of galactose and copper with *C. neoformans* metabolism and virulence, and difficulty regulating expression in animal hosts limit their utility in studying essential genes both *in vitro* and *in vivo* (11-13).

Compared to other regulatable expression systems, the tetracycline-repressible promoter system has superior expression control and compatibility in laboratory settings and infection models. This regulatable promoter system has been successfully adapted in *S. cerevisiae* and other fungal pathogens such as *C. albicans* and *Candida auris* (3, 14, 15). Typically, this system has two components: 1) a chimeric transactivator protein consisting of the *Escherichia coli tet*R DNA-binding domain fused to transcriptional activation domains, and 2) a tetracycline response element (TRE) containing multiple repeats of tetracycline operator sequence to which the transactivator binds (14). Binding of the transactivator protein to TRE enables constitutive gene expression and addition of tetracycline (and analogues such as doxycycline (DOX)) blocks association between the transactivator and TRE resulting in repression of a Tet-promoter regulated gene (Figure 1A). Importantly, the addition of DOX can reach a steady state above 1 µg/mL in mouse plasma and major organs by feeding animals with DOX-supplemented drinking water or chow, enabling the study of gene essentiality *in vivo* (4, 16-18). Despite superior *in vitro* and *in vivo* expression regulation, the tetracycline-repressible system has yet to be adapted in *C. neoformans*.

**Figure 1.**
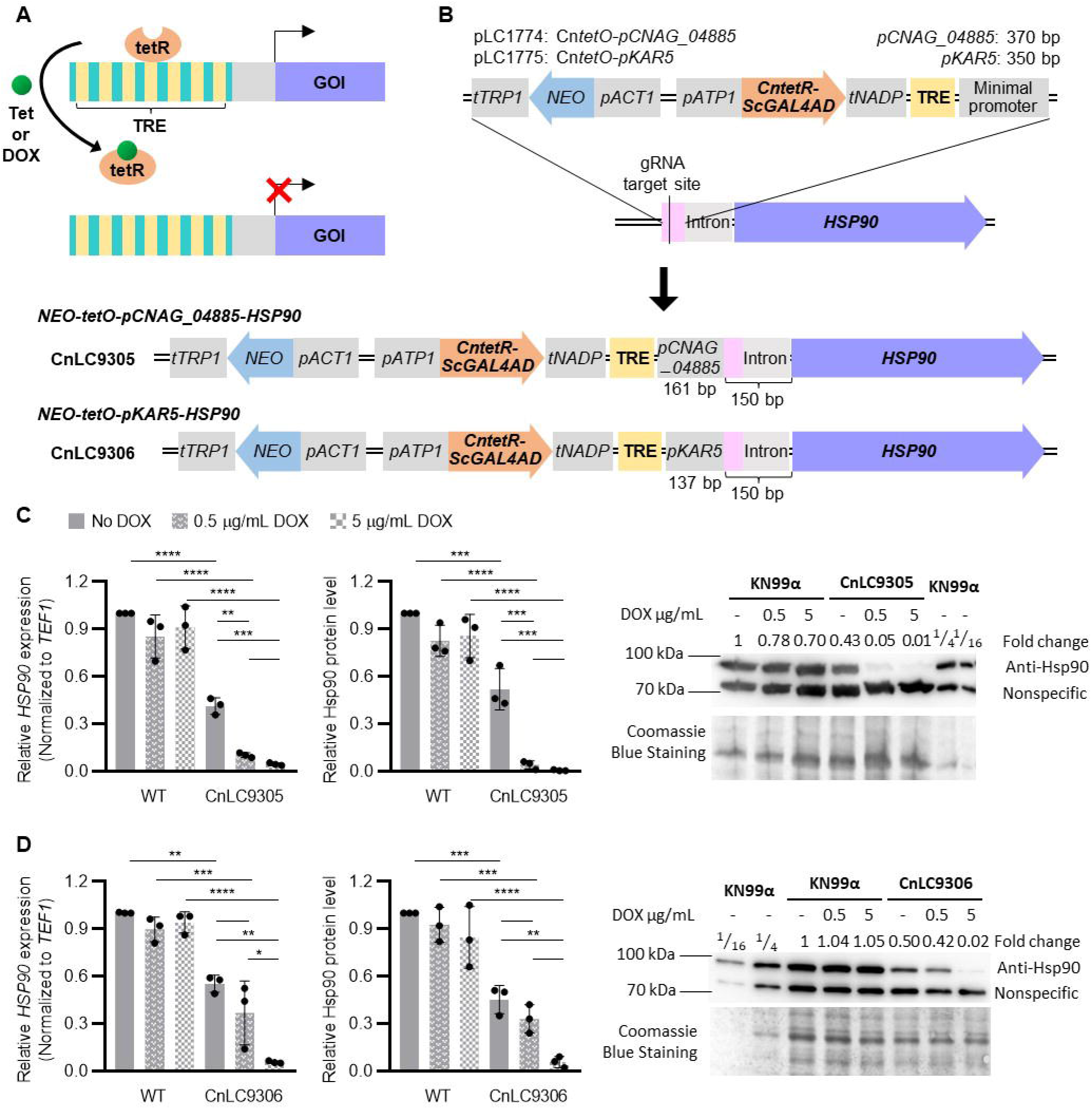
Adaptation of tetracycline-repressible system in *Cryptococcus neoformans*. **A**. Schematic diagram for the tetracycline-repressible system. In the presence of tetracycline (Tet) or doxycycline (DOX), gene expression is repressed due to disassociation of the transactivator (tetR) from the tetracycline response elements. **B**. Schematic diagram of the tetracycline-repressible system in *C. neoformans* for the generation of two *tetO-HSP90* strains. A *C. neoformans* G418 selection marker under the control of the *ACT1* promoter, the *C. neoformans* codon optimized transactivator (tetR) with the activation domain from *S. cerevisiae GAL4* gene, the tetracycline response element (TRE) containing seven repeats of tetracycline operator sequence, and minimal promoter from either *CNAG_04885* or *KAR5* were assembled into two *tetO* vectors, pLC1774 and pLC1775. *CntetO* cassettes were integrated in the promoter region of *HSP90* via CRISPR/Cas9 genome editing. Doxycycline-sensitive *tetO-HSP90* strains CnLC9305 and CnLC9306 were identified, and Sanger sequencing confirmed the integration of the tetO cassettes 150 bps upstream of the *HSP90* start codon. The minimal promoters *pCNAG_04885* and *pKAR5* were truncated to 161 bps and 137 bps, respectively. **C-D)** Wild type (WT, KN99_α_) and the two *tetO-HSP90* strains, CnLC9305 and CnLC9306, were grown in liquid YPD media in the presence or the absence of 0.5 µg/mL DOX overnight. On the second day, YPD cultures were sub-cultured back into YPD, and YPD with DOX cultures were sub-cultured into YPD containing 0.5 or 5 µg/mL DOX for four to five hours at 30 °C. Cells were harvested to examine *HSP90* expression levels by real time-PCR and Hsp90 protein levels by western blot for (**C**) CnLC9305 and (**D**) CnLC9306. Relative expression levels for *HSP90* were normalized to *TEF1* and compared to wild-type cells grown in YPD medium. Each datapoint represents the mean of technical triplicates, normalized to the *HSP90* expression level in wild-type cells grown in YPD medium. Relative Hsp90 protein levels were normalized to the nonspecific protein band recognized by the anti-Hsp90 antibody and compared to wild type grown in YPD medium. Representative western blot image from one biological experiment for each strain is provided to visualize Hsp90 protein levels. Wild type (KN99_α_) protein sample was diluted four-fold and 16-fold to visualize the decrease in protein band intensity. Hsp90 protein levels were normalized to nonspecific protein bands recognized by the anti-Hsp90 antibody and compared to wild type grown in YPD medium. Hsp90 protein level fold changes are listed above the western blot image and Coomassie stain of the membrane confirming equal loading is shown below, which matches the pattern observed with the nonspecific band. Error bars in bar graphs represent standard deviation of the mean for biological triplicates. Two-way ANOVA with Bonferroni’s correction was performed to determine statistical significance. ^*^ *p* ≤ 0.05, ^**^ *p* ≤ 0.01, ^***^ *p* ≤ 0.001, ^****^ *p* ≤ 0.0001.

In this observation, we describe adaptation of the tetracycline-repressible system in *C. neoformans* for modulating essential gene expression, focusing on the well-characterized essential gene *HSP90* as proof of concept (Figure 1B) (12, 19). To do so, we first codon-optimized the tetracycline-responsive transactivator (tetR) with the Gal4 activation domain from *S. cerevisiae* based on *C. neoformans* codon usage (Figure S1) (3, 20). This sequence was then cloned into vectors that included the tetracycline operator (*tetO*) cassettes with a G418-selectable drug marker, *C. neoformans* codon optimized *CntetR-ScGAL4AD*, tetracycline response element (TRE), and minimal promoters from either *CNAG_04885* (to generate CnLC9305) or *KAR5* (to generate CnLC9306) for efficient transcription initiation in *C. neoformans* (Figure 1B). Minimal promoters were selected based on their low expression profiles from published transcriptomic profiling studies to avoid potential competing regulation with the tetracycline-repressible system (21, 22).

The assembled *tetO* cassettes were introduced into wild-type KN99_α_via the short homology-directed CRISPR/Cas9 genome editing technique to replace 111 bps upstream of the first *HSP90* intron (pink box in Figure 1B) (20, 23). This location was selected to maintain the integrity of the intergenic intron upstream of *HSP90* start codon that is likely required for proper expression. Transformants were phenotypically screened for DOX-dependent growth phenotypes and one candidate for each *tetO* cassette was selected for downstream characterization (Figure 1B). Interestingly, Sanger sequencing of the prioritized transformants showed that only 17 bps containing the Cas9 recognition PAM site in the targeted 111 bp-sequence were replaced by *tetO* cassettes. Additionally, both minimal promoters were truncated from 370 to 161 bps for *pCNAG_04855* and from 350 to 137 bps for *pKAR5*. The non-homologous integration occurring downstream of the Cas9-mediated double-strand break was likely facilitated by the NHEJ pathway (Figure 1B).

To characterize the activity of the *tetO* cassettes in *C. neoformans*, the two *tetO-HSP90* strains, CnLC9305 and CnLC9306, were examined for *HSP90* transcript and protein levels in the absence and presence of DOX (Figure 1C and 1D). In the absence of DOX, both strains showed significantly reduced *HSP90* transcript and protein levels, which were about half of the wild-type levels. In the presence of 5 µg/mL (∼11.2 µM) DOX, these levels were significantly depleted to below 6% of wild-type levels. At a lower DOX concentration of 0.5 µg/mL (∼1.12µM), *HSP90* mRNA and protein levels were significantly depleted in CnLC9305, but not in CnLC9306, indicating a greater sensitivity to DOX with the truncated minimal promoter *pCNAG_04885* (Figure 1C and 1D).

To visualize the impact of DOX treatment on *C. neoformans* growth, wild-type and the *tetO-HSP90* strains were cultured on both solid YPD agar medium and liquid YPD medium with and without DOX supplementation (Figure 2). Specifically, on YPD agar, 5 µg/mL DOX significantly reduced growth of CnLC9305 and CnLC9306 at 30 °C and completely blocked growth at 37 °C. In agreement with the observed DOX regulation of *HSP90* at the transcript and protein levels, 0.5 µg/mL DOX had a more dramatic effect on growth of CnLC9305 at 37 °C compared to CnLC9306 (Figure 2A). Interestingly, despite the reduced expression levels of *HSP90* in the absence of DOX for the *tetO-HSP90* strains, minimal to no growth inhibition was observed for CnLC9305 and CnLC9306 when compared to wild type at either temperature (Figure 2A and 2B). Similar results were observed in liquid medium with levels of DOX below 1 µg/mL reducing growth and metabolic activity of both strains, and CnLC9305 showing increased sensitivity to DOX. Finally, we cultured *tetO-HSP90* strains in the absence and presence of a high concentration of DOX (20 µg/mL) over four days and measured viability by quantifying colony forming units (CFUs) of the surviving populations. Wild-type cells treated with the pharmacological Hsp90 inhibitor radicicol (RAD) at an inhibitory concentration of 10 µM was used as a control. Similar to RAD treatment, DOX reduced viability of both *tetO-HSP90* strains, with a two-log reduction in CFU compared to wild type and no DOX controls after 72 hours. Overall, this characterization confirms the successful generation of conditional-expression strains that regulate essential genes in *C. neoformans* and highlights that the selection of a minimal promoter in the *tetO* cassette plays a significant role in modulating expression of downstream genes.

**Figure 2.**
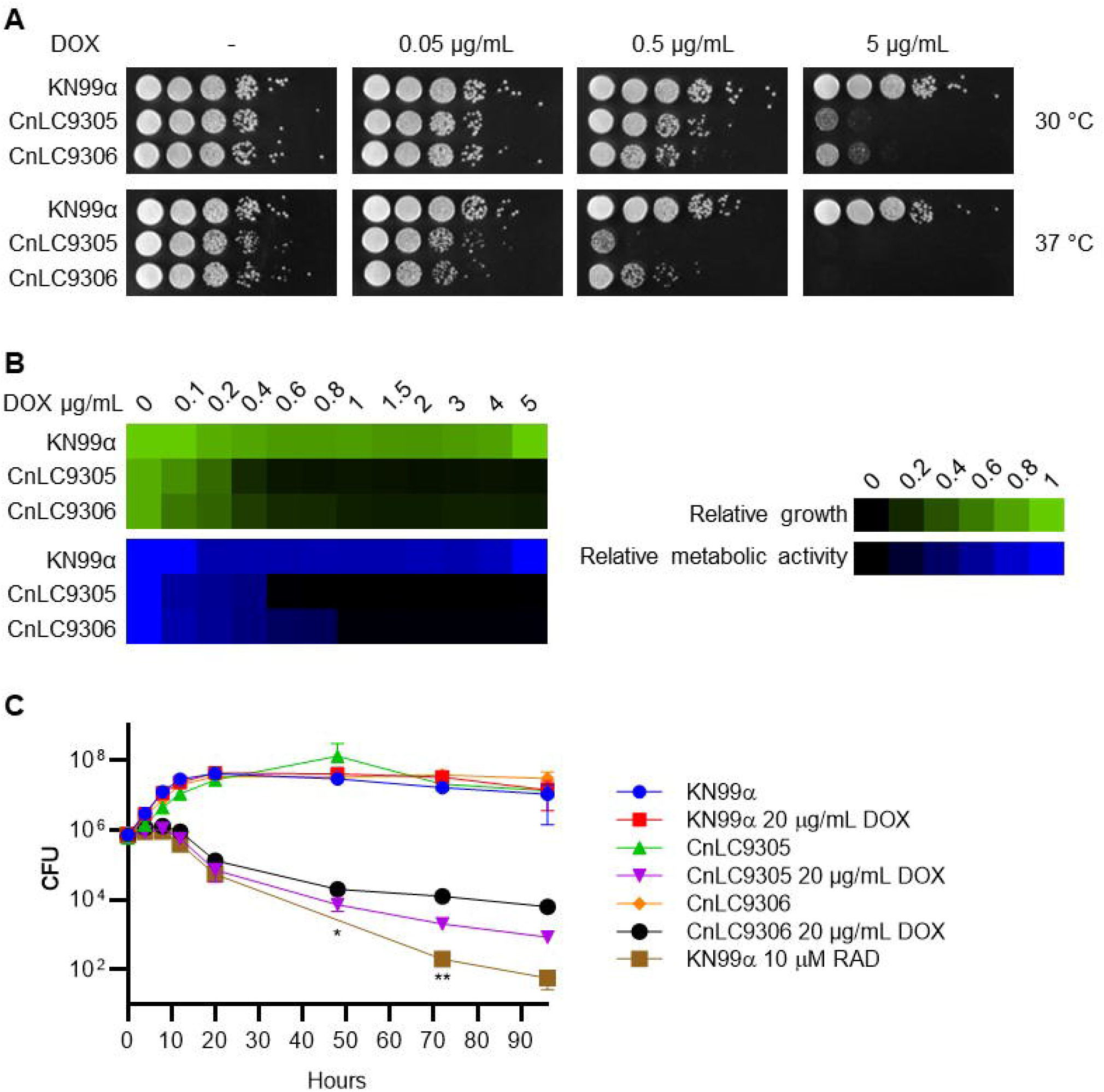
Characterization of doxycycline sensitivity in the *tetO-HSP90* strains. **A**. To determine DOX sensitivity on solid YPD agar medium, cells from wild-type, CnLC9305, and CnLC9306 overnight cultures in liquid YPD medium were 10-fold serially diluted at a starting optical density (OD_600_) of 0.8 and spotted on YPD agar medium. Cells from overnight cultures in liquid YPD medium with 0.5 µg/mL DOX were similarly diluted and spotted on YPD agar medium supplemented with 0.05, 0.5, or 5 µg/mL DOX. Agar plates were incubated at 30 °C or 37 °C and imaged after two days. **B**. To determine DOX sensitivity in liquid YPD medium, cells from wild-type, CnLC9305, and CnLC9306 overnight cultures in liquid YPD medium were washed and cell densities were determined using a hemocytometer. For each strain, 50,000 cells were seeded in 200 µL liquid YPD medium with DOX at a gradient of concentrations ranging from 0 to 5 µg/mL in a 96-well plate and the plate was incubated at 37 °C for 72 hours. OD_600_ was measured using a plate reader and relative growth was normalized to wild type without DOX. To measure metabolic activity, Alamar blue was added to each well at a dilution factor of 1:40 and the plate was incubated in the dark at room temperature for 12 hours before fluorescence measurement, with an excitation wavelength of 535 nm and an emission wavelength of 595 nm. Relative metabolic activity was normalized to wild type without DOX. **C**. To determine viability of the *tetO-HSP90* strains, wild type, CnLC9305, and CnLC9306 were inoculated at 1 × 10^6^ CFU in liquid YPD medium with or without 20 µg/mL DOX. Wild type grown in liquid YPD medium supplemented with 10 µM radicicol was used as a control. Cells were grown at 37 °C under shaking condition at 220 rpm, and plated on YPD agar medium at 0, 4, 8, 12, 20, 48, 72, and 96 hours post inoculation to determine CFU. Error bars represent standard deviation of the mean for biological triplicates. All three data points (^*^) and two datapoints (^**^) were below the detection limit of 2000 CFU for wild type grown in the presence of 10 µM radicicol.

Compared to our previous use of the copper-repressible *pCTR4* promoter (12), the tetracycline-repressible system presented here displays superior regulation of the essential gene *HSP90*, which suggests the adapted *tetO* system is a powerful tool to study essential gene function in *C. neoformans*. Similarly, an auxin-inducible degron system has recently been developed to control essential gene expression in *C. neoformans* with promising results (24). Furthermore, in the recent study by Peterson et al., we describe the generation of a marker-free, diploid wild-type *C. neoformans* strain CnLC6683, a fusion product of the congenic wild-type strain pair KN99_α_and KN99**a**, which offers more flexibility in genome editing and enables haploinsufficiency studies in this typically haploid fungal pathogen (25). Combination of this diploid strain and the newly adapted tetracycline-repressible system promises a fruitful exploration of essential gene functions in *C. neoformans* and improves our ability to perform functional genomic analyses in this important human fungal pathogen.

## Acknowledgements

We thank ZhongLe Liu, Saif Hossain, Luke Whitesell, and Emily Xiong for providing critical suggestions on the manuscript. We also thank the Cowen lab members for thoughtful discussions.

L.E.C. is supported by a National Institutes of Health (NIH) R01 grant (R01AI165466), and is a Canada Research Chair (Tier 1) in Microbial Genomics & Infectious Disease. LEC is the co-director of the CIFAR Fungal Kingdom: Threats & Opportunities program.

C. F. conceived the study, performed all experiments, interpretated data and results, and wrote and edited the manuscript. N.R. and L.E.C. interpreted the data and results and helped write and edit the manuscript.

---

**Figure S1.**
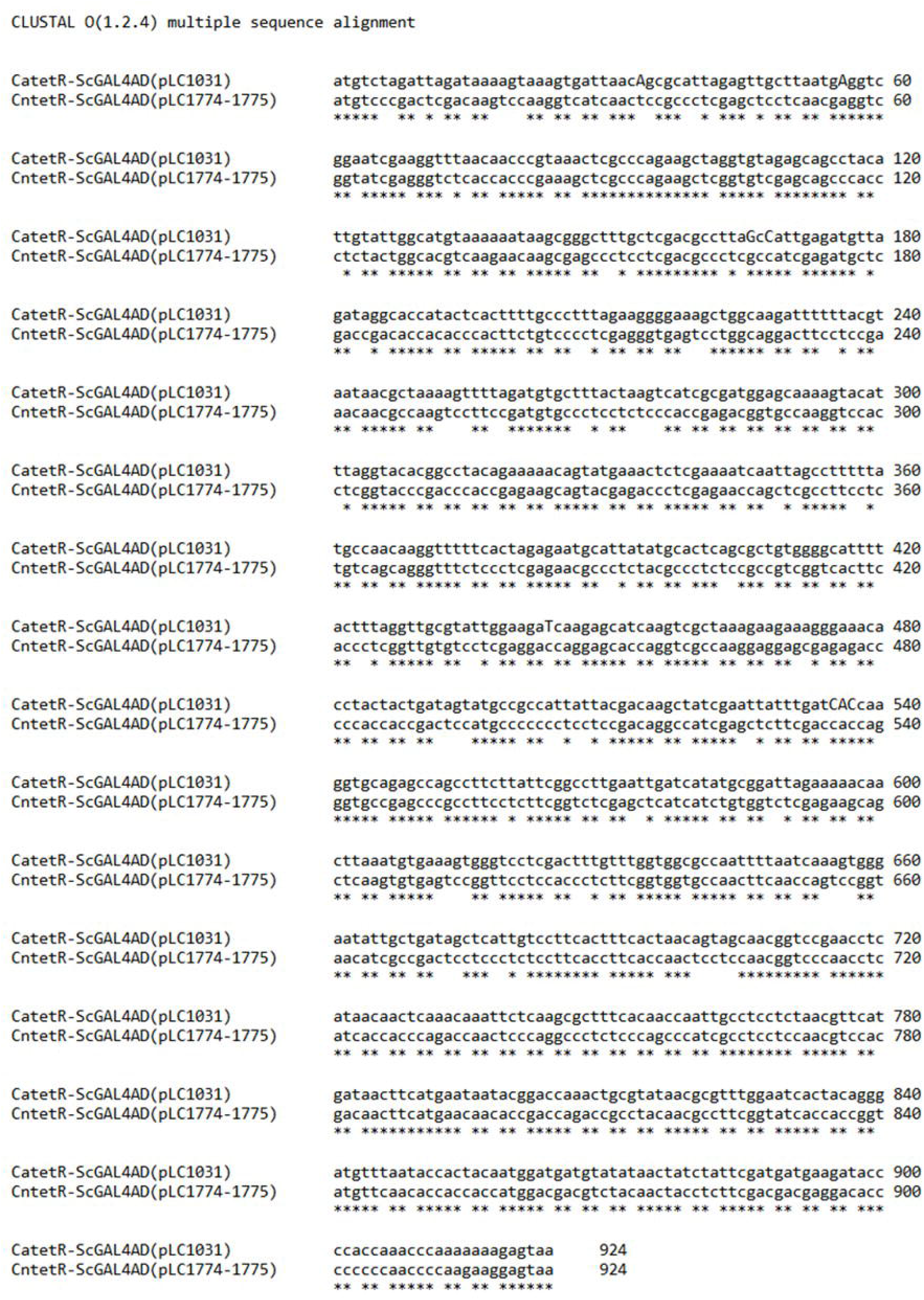
Sequence alignment of codon-optimized *CntetR-ScGAL4AD* with *CatetR-ScGAL4AD*. Clustal alignment was performed on the sequences of codon-optimized *CntetR-ScGAL4AD* and *CatetR-ScGAL4AD* to highlight the changes in nucleotide sequence.

